# Epistasis Detection using Model Based Multifactor Dimensionality Reduction in Structured Populations

**DOI:** 10.1101/541946

**Authors:** Fentaw Abegaz, François Van Lishout, Jestinah M Mahachie John, Kridsadakorn Chiachoompu, Archana Bhardwaj, Elena S. Gusareva, Zhi Wei, Hakon Hakonarson, Kristel Van Steen

## Abstract

In genome-wide association studies, the extent and impact of confounding due population structure have been well recognized. Inadequate handling of such confounding is likely to lead to spurious associations, hampering replication and the identification of causal variants. Several strategies have been developed for protecting associations against confounding, the most popular one is based on Principal Component Analysis. In contrast, the extent and impact of confounding due to population structure in gene-gene interaction association epistasis studies are much less investigated and understood. In particular, the role of non-linear genetic population substructure in epistasis detection is largely under-investigated, especially outside a regression framework. In order to identify causal variants in synergy, to improve interpretability and replicability of epistasis results, we introduce three strategies based on model-based multifactor dimensionality reduction (MB-MDR) approach for structured populations. We demonstrate through extensive simulation studies the effect of various degrees of genetic population structure and relatedness on epistasis detection and propose appropriate remedial measures based on linear and non-linear sample genetic similarity.

**Authors Summary:** One of the biggest challenges in human genetics is to understand the genetic basis of complex diseases such as cancer, diabetes, heart disease, depression, asthma, inflammatory bowel disease and hypertension, for instance via identifying genes, gene-gene and gene-environment interactions in association studies. Over the years, a more prominent role has been given to gene-gene interaction (epistasis) detection, in view of precision medicine and the hunt for novel drug targets and biomarkers. However, the increasing number of consortium-based epistasis studies that are marked by heterogeneous sample collections due to population structure or shared genetic ancestry are likely to be prone to spurious association and low power detection of associated or causal genes. In this work we introduced various strategies in epistasis studies with correction for confounding due to population structure. Based on extensive simulation studies we demonstrated the effect of genetic population structure on epistasis detection and investigated remedial measures to confounding by linear and nonlinear sample genetic similarity.

## Introduction

Genome-wide association studies (GWAS) are an effective approach for identifying genetic variants associated to disease risk^1^. In the context of such studies, population stratification refers to a systematic ancestry differences between cases and controls^1^. The phenomenon is of particular concern in study designs with unrelated individuals. In contrast, family-based genetic association studies offer protection from population stratification, by using family data as internal controls, although at the expense of some loss of power from genotypic overmatching ^2,3^. For case-control genetic association studies, spurious associations are caused by the co-occurrence of two factors: a difference in proportion of individuals from two (or more) subpopulations in cases and controls, and subpopulations having differing allele frequencies at the locus under investigation. This is in fact a special case of Simpson’s Paradox ^4^. In general, this statistical phenomenon causes a potential bias in data analysis and occurs when a relationship or association between two variables reverses when a third factor, called a confounding variable, is introduced. The paradox also occurs if an association reverses when the data is aggregated over a confounding variable. Increasing the sample size is usually not a remedy for this issue, but may worsen the problem ^5^. Several causes exist for population stratification. The basic one being shared genetic ancestry as a result of non-random mating between subgroups in a population due to various reasons, which may include social, cultural or geographical ones. From an evolutionary point of view, not only population stratification, but also admixture (i.e., inter-mating between genetically distinct groups) is created by human mating patterns. Potential consequences of population stratification are confounding, cryptic relatedness (i.e., unobserved ancestral relationships between individual cases and controls causing them to be non-independent) and selection bias ^6,7^.

In case/control GWA studies, several strategies have been introduced in the literature for protecting against population structure mainly based on Principal Components Analysis (PCA). In contrast, the extent and impact of confounding due to population structure in gene-gene interaction studies is much less investigated and understood. However, the growing interest in the importance of detecting gene–gene interactions in the development and progression of complex diseases has led to the development of several tools; to name but a few: generalized linear regression models (GLM), BOOST ^8^, Model Based Multifactor Dimensionality Reduction (MB-MDR) ^9,10^, Multifactor Dimensionality Reduction (MDR) ^11^, Random Forest ^12^, PLINK ^13^, BiForce ^14^, Bayesian Models (e.g., BEAM) ^15^ and several others. For extensive reviews and appropriate references, please refer to ^16–20^. However, the literature on epistasis detection in structured populations is very limited, apart from scenarios using a regression framework for association testing. On the other hand, Model-Based Multifactor Dimensionality Reduction (MB-MDR) offers a general framework and software tool for epistasis detection that can offer flexible maneuvering between different measurement scales for phenotypes and genomic predictors ^9,10,21^. Interestingly, the MDR-SP method ^22^, to our knowledge the only MDR-inspired method that can deal with structured populations, combines MDR ^11^ with ideas implemented in the EIGENSTRAT software ^23^.

In this article, we introduce strategies to account for population structure in epistasis studies using the MB-MDR framework. In particular, for the remainder of this article, we restrict attention to case-control study designs (binary original traits) and biallelic Single Nucleotide Polymorphisms (SNPs) as genetic markers. We propose and fully describe three strategies: i) MBMDR-PC, ii) MBMDR-PG and iii) MBMDR-GC. In MBMDR-PC, principal components (PCs) adjusted phenotypes but original genotypes are used to detect epistatic SNP pairs, similar to ^23^. In MBMDR-PG adjusted phenotypes are obtained from fitting logistic mixed (polygenic) models on the original binary trait, hereby allowing to adjust for additional structure such as those arising from family relationships and cryptic relatedness. In MBMDR-GC, we follow principles of Genomic Control correction in GWAS, but allow for multi-locus adaptivity. These methods are evaluated via extensive simulation studies which, to our knowledge, are unique in their kind in that complex non-linear population structures structural epistasis are considered as well. Here, we let structural epistasis refer to the presence of interacting markers driving population differences or population substructure. All proposed strategies are formally compared to MDR-SP ^22^, in terms of type I error control and statistical power. Our work is important to highlight the impact of nonlinear genetic population substructure in epistasis detection.

## Material and Methods

All proposed genome-wide epistasis screening strategies in structured populations are built on the Model-Based Multifactor Dimensionality Reduction method (MB-MDR) ^9,10,24,25^. Even though the MB-MDR framework can be used for higher level interaction detection and various outcome measurement scales and study designs, here we restrict attention to pair-wise interactions. A graphical overview of the newly introduced methods is provided in Figure 1 and explained in more detail below.

**Figure 1.**
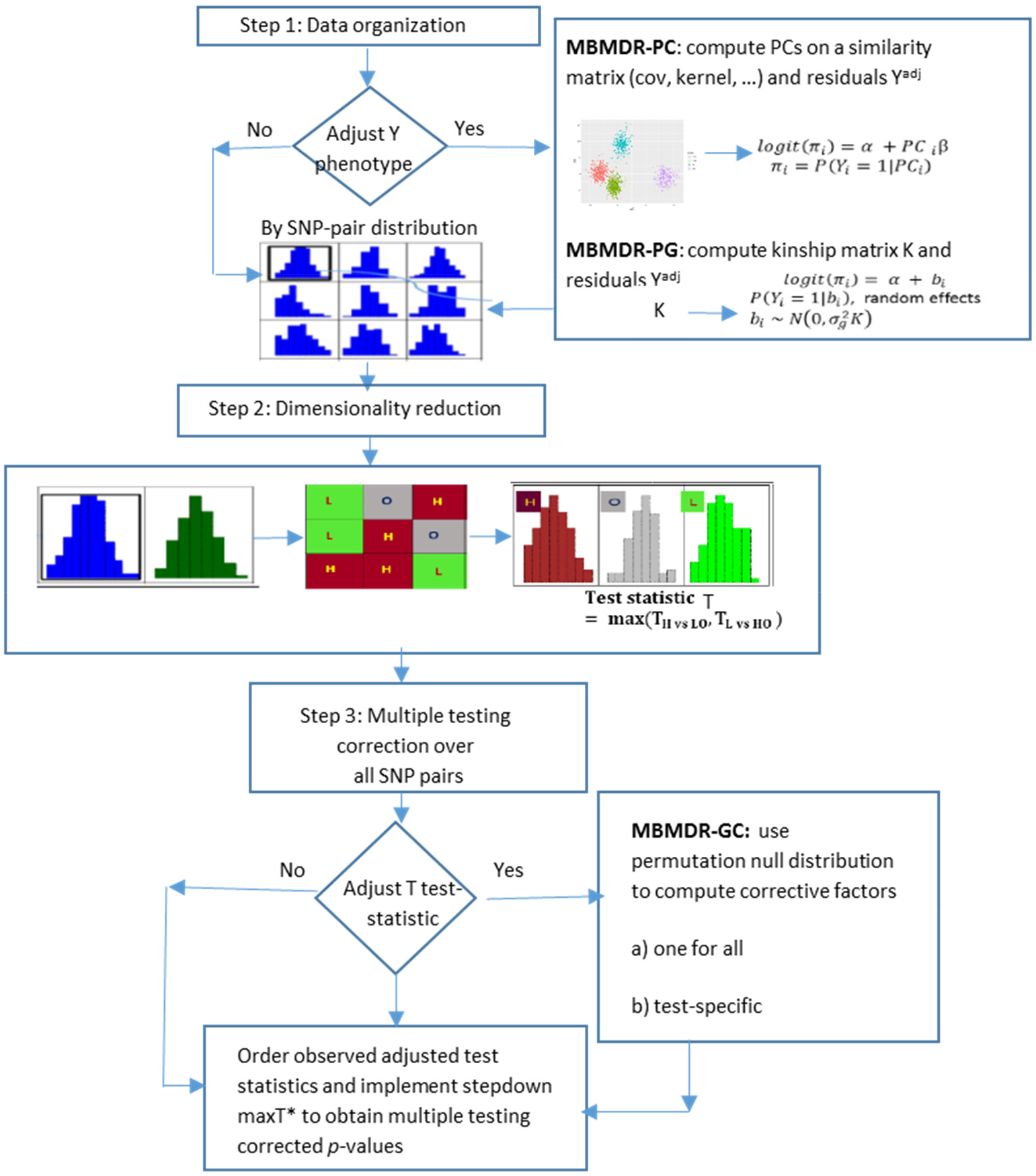
Extended graphical presentation of the key steps involved in MB-MDR for structured populations

### MBMDR-PC: accounting for genomic structure by PCs

Similar to EIGENSTRAT ^23^, we use either linear or non-linear (kernel) PCs to correct for population structure, and coin our strategy MBMDR-PC. The adopted procedures to compute linear and non-linear PCs via genome data are outlined in the Supplementary Material.

In MBMDR-PC we append the classic data organization step in MB-MDR with one that computes new phenotypes, adjusted for population structure. The new phenotypes are taken as input to classic MB-MDR, in an attempt to capture genetic interactions that are not spurious due to inadequate handling of population structures. In what follows, we give a detailed outline of MBMDR-PC (see also Figure 1).

#### Step 1: Data organization

As in classic MB-MDR, for every two SNP loci *j* and *k*, individuals with non-missing genotype data will exhibit one of nine possible two-locus genotypes {00, 01, 02, 10, 11, 12, 20, 21, 22}, for an individual’s genotype *G_j_* at locus *j* and *G_k_* at locus *k* taking on 0,1,2 as possible values (i.e., number of minor alleles). Specific to MBMDR-PC, the regression framework is adhered to create new phenotypes that are adjusted for population structure captured by a number of PCs. In case-control epistasis studies, where the phenotype *Y* represents disease status (1 affected, 0 unaffected), and *G_j_* and *G_k_* refer to genetic information at two SNP loci *j* and *k* (for example using additive encoding), epistasis can be investigated by making inference on the interaction parameter *θ* in the following logistic regression model

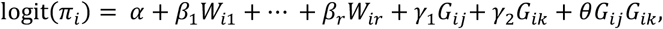

where *π_i_* = *P*(*Y_i_* = 1 | *W*_*i*1_, …,*W_ir_, G_ij_, G_ik_*) is the probability of disease for subject *i*, possibly conditional on the first *r* principal components *W*_*i*1_, … *W_ir_*. The vector *β* = (*β*_1_,…,*β_r_*) is a vector of regression parameters corresponding to the r principal components, *α* is the intercept term, and *γ*_1_ and *γ*_2_ are the main effects of the two SNPs and *θ* captures the interaction effect between the two SNPs at loci *j* and *k*. Earlier reports of limitations related to logistic regression for higher-order interaction modeling ^26^, including having to make “model assumptions” about mode of inheritance (i.e., related to choosing a particular encoding scheme for genetic exposures) lies at the basis of Model-Based Multifactor Dimensionality Reduction (MB-MDR), which is non-parametric in its core. However, when adjustments need to be made for lower order effects or confounders, the MB-MDR paradigm needs to be combined with the regression paradigm. Related to MBMDR-PC, we derive adjusted phenotypes from the aforementioned logistic regression model by subtracting model-fitted values from observed phenotype values:

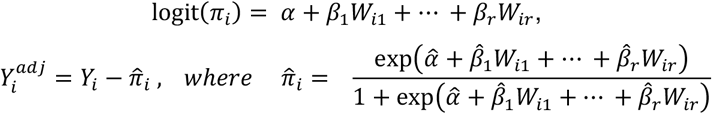

This can be accomplished in R using the package ***glm*** (Generalized Linear Models) ^27^. The fitted model should be adapted according to the measurement type of the original phenotype, by selecting an appropriate link function linking the linear predictor *α* + *β*_1_*W*_*i*1_ + ⋯ + *β_r_W_ir_* to the (adjusted) phenotype. Conditioning on additional confounders (e.g., sex, age, etc.) is straightforward by including them in the expression for *logit*(*π_i_*) above.

#### Step 2: Multi-locus prioritization and dimensionality reduction

The adjusted phenotype obtained in *Step 1* is subsequently investigated for distributional differences between multilocus genotypes (*G_j_* × *G_k_*). For SNPs at loci *j* and *k*, such investigations are reduced to making inferences about *γ* as in

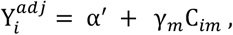

for which 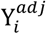 denotes the adjusted phenotype (*Step 1*) and *C_im_* is 1 if the m-th multilocus genotype derived from *G_j_* and *G_k_* is 1 and 0 otherwise. In practice with MBMDR, a Student’s t-test is carried out, for each multilocus genotype cell C_m_ at the liberal significance level of 0.10, comparing the mean of cell C_m_ with the mean of the remaining eight cells. If the test is not significant at 0.10 level, the cell is labeled as O (no evidence for risk). If the test is significant at 0.10 the cell is labeled as H (high risk) or L (low risk). The sign of the test statistic is used to distinguish between H and L: a positive (negative) sign refers to risk H (L). The thresholding of 0.10 is motivated in earlier work ^24^.

The result is thus a new categorical variable with values H, L and O, which captures information about the importance of the pair of SNPs with respect to the adjusted phenotype. Subsequently, a final aggregate association test is performed on the new construct and the adjusted phenotype. In particular, we consider the maximum of two t-test statistics denoted by *T_H_* and *T_L_*, again comparing the mean responses of H versus {L, O} categorized individuals and the means of L versus {H, O} labelled individuals, respectively. Contrast tests, comparing H versus L combined multilocus genotypes per SNP pair can be considered as well. These have been evaluated elsewhere ^24^ but were not considered for the purposes of this paper.

#### Step 3: Significance assessment

For every pair of SNPs in the data we obtain a single test statistic given by

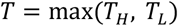

from Step 2. This maximum test statistic will no longer follow a t-distribution, due to the compounding of evidences in Step 2. The significance of *T* per SNP pair is therefore assessed via resampling based strategies. In particular, here we generate 999 permutation-based replicates by permuting adjusted trait labels, yet keeping the correlation structure between SNPs intact. Multiple testing is taken care of by a step-down maxT approach as described in-adjusted p-values ^28^, hereby ensuring partial strong control of FWER at 5%. Strong control properties can be stated under the assumption of the subset pivotality property ^29^. For large samples, such as in the real-life data application, we adhered to an approximated step-down maxT adjusted p-values, as described in ^28^.

### MBMDR-PG: accounting for genomic structure due to families and cryptic relatedness via the extended Polygenic Model

Family structure or cryptic relatedness may induce phenotypic similarity between individuals and may confound gene-phenotype associations in GWAS when not properly accounted for. Whereas PCs have proven useful in GWAs and structured populations due to shared genetic ancestry, they are not suitable to adequately protect for the effects of familial or cryptic relatedness on GWAS ^1^. With the recent developments of computationally efficient algorithms, mixed models have become feasible in the context of GWAS as well as GWAIS, in structured populations, whether this structure presents population stratification, known or unknown relatedness. For quite some time, GWAS for binary traits have been analyzed with linear mixed models, assuming that little harm is done when sample sizes are in the thousands as is often the case with consortium data ^30^. However, Chen et al. ^31^ showed that linear mixed models are really inappropriate for analyzing binary traits when population stratification induces violation of the constant residual variance assumption in linear mixed models. Therefore, these authors developed a computationally efficient logistic mixed model for binary trait GWAS in the presence of population structure as well as familial and cryptic relatedness. In the same spirit of the logistic regression models adopted before, a logistic mixed model that includes interaction effect between two SNPs can be defined as

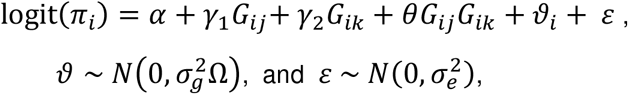

where *π_i_* = *P*(*Y_i_* = 1|*G_ij_, G_ik_*, ϑ) is the probability of disease for subject *i*, conditional on SNPs and random effects *ϑ_i_*. Here, *ϑ* is a *N* × 1 vector of random effects assumed to follow a multivariate Gaussian distribution and 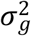 is the additive genetic variance Ω is the genetic similarity matrix between all pairs of individuals (dimension *N* × *N*) such that Ω_*il*_ represents the similarity between individuals *i* and *l*. An estimate of the genetic similarity matrix, Ω, is required which can be obtained from a large number of genetic variants ^32^. Fitting the model involves integrating over the random effects vector B with respect to the Gaussian distribution so that the likelihood is maximized with respect to the parameters 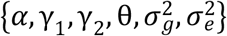 ^33^. In MBMDR-PG we obtain the adjusted phenotype from the residuals of fitting the logistic random effect model using the R package ***GMMAT*** (Generalized Linear Mixed Model Association Test) ^31^. Then, similar to MBMDR-PC we use the adjusted phenotype as input for interaction analysis with MBMDR as described above in *Steps 2-3*.

### MBMDR-GC: accounting for genomic structure via genomic control

The genomic control method introduced in ^34^ is computationally simple and fast to control for population structure in case-control association studies. The key idea is to divide the observed association test statistic by a single factor, *λ_GC_*, which measures the overall inflation in association test statistic due to population stratification. The factor *λ_GC_* can be estimated by dividing the mediums of the observed association test statistics across a set of markers by the theoretical median of the association test statistic. Notably, corrective factors computed in this sense may turn out to be less than 1 and may actually inflate observed test values rather than deflating them. Although genomic control have proven useful in a variety of contexts, Price et al. ^23^ pointed out that the common deflation factor applied to all SNPs where some SNPs differ in their allele frequencies across ancestral populations more than others could lead to loss of power. As a solution ^35^ considered test specific genomic control. MBMBDR-GC also employs test-specific genomic control, adapted to the MB-MDR testing framework.

In MBMBDR-GC principles of classic GC in GWAS for structured populations are adopted ^23^. Large differences between several multi-locus genotype frequencies across populations may lead to power loss when a single corrective inflation factor GC is used. Therefore, in MBMDR-GC the definition of GC is adapted and the permutation null data generated in MBMDR (*step 3*, Figure 1) is exploited to estimate a test-specific GC factor, similar to ^35^. In particular, the j^th^ SNP-SNP interaction pair, the corrective factor *λ_GC,j_* is estimated as

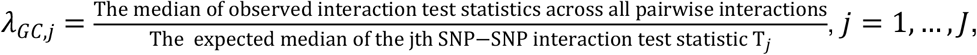

where *J* is the total number of pairwise interactions and for which the expected median of the j^th^ SNP-SNP interaction test statistic is computed from 1000 permutations under the null distribution constructed by randomly permuting the phenotype values. Then the adjusted test statistic for the j^th^ interaction pair becomes *T_j_*/*λ_GC,j_*.

### Application on synthetic data

To assess type I error control and power performance of MBMDR-PC, MBMDR-PG and MBMDR-GC, and to compare it to MDR-SP ^22^, we set up a series of simulation settings, involving either unrelated or related individuals, as depicted in Figure 2. Two main strategies were adopted for our simulation study with unrelated individuals involving: 1) HapMap data as a template with and without structural epistasis, 2) newly generated discrete populations without a reference template, with and without structural epistasis (referred to as model-based data). More detailed explanations are given next.

**Figure 2.**
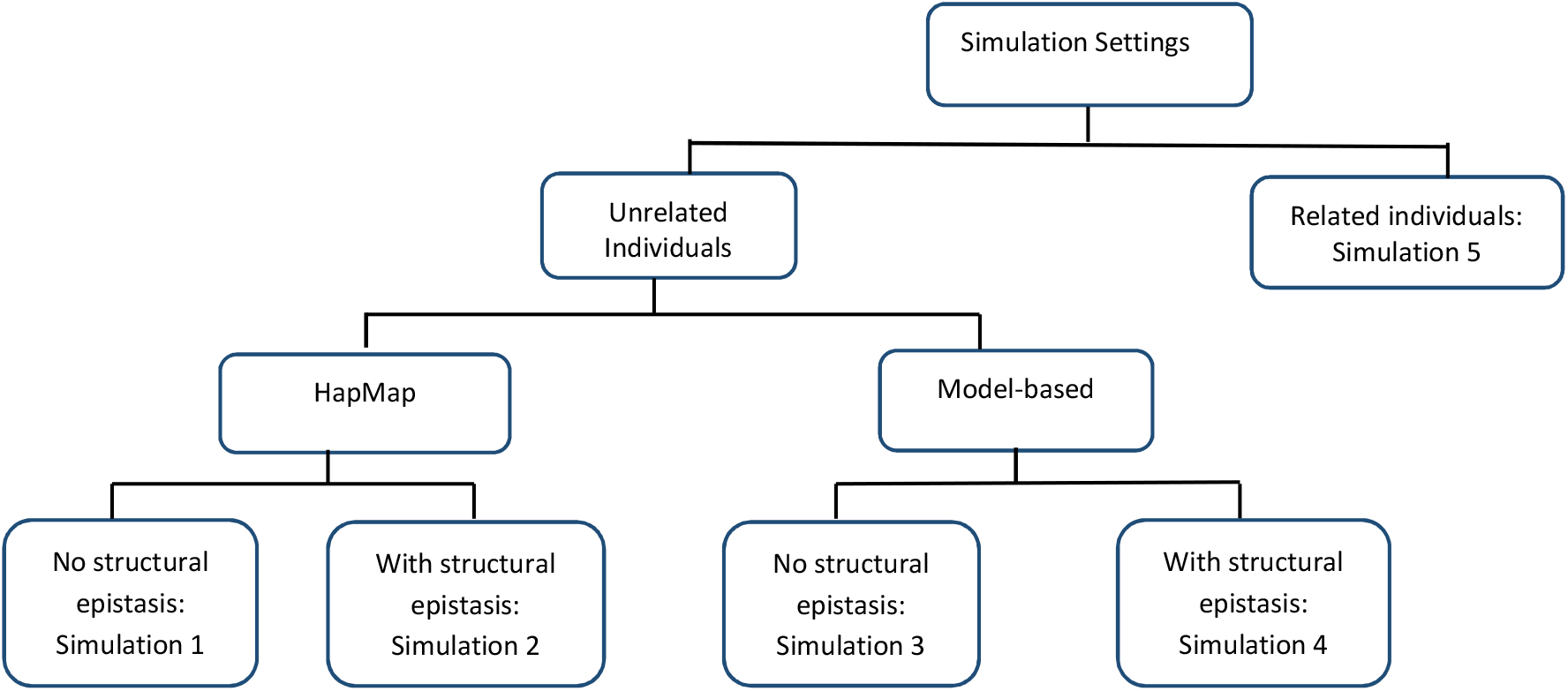
Flow of considered simulation settings.

In summarizing the simulation results, type I error rates are obtained as the proportion of the number of simulated datasets for which a pair of SNPs found significant at the 5% level after correcting for multiple testing. Similarly, the power is obtained as the proportion of the number of simulated datasets for which only the functional pair of SNPs found significant at the 5% level after correcting for multiple testing.

#### ***Simulation setting 1***: *Synthetic* data derived from HapMap in the absence of structural epistasis

For each simulated set, we considered 200, 300 or 400 individuals, half of them cases and half of them controls, 80% controls and 40% cases as European (CEU) and 20% controls and 60% cases as African (YRI). We followed the simulation strategy adopted for MDR-SP ^22^ to generate genotype data with unlinked SNPs. In particular, *L* ∈ {200, 400, 800}, independent SNPs were randomly selected from the total number of SNPs from the pooled HapMap3 (CEU and YRI) data with quality control that includes (only founders, HWE *p*-value threshold of 0.001, individual and genotype missing rates of 5% and 2%,respectively, minor allele frequency *MAF* > 0.05 and LD pruning threshold of *r*^2^ = 0.75) (http://www.sanger.ac.uk/resources/downloads/human/hapmap3.html), and minor allele frequencies were extracted for these two populations. Genotypes were then generated under the assumption of Hardy-Weinberg equilibrium (HWE). The genotypes for the *L* unlinked SNPs were subsequently used to compute principal components and the first 10 principal were retained to capture population substructure.

Since the aim of this study is not to evaluate multiple testing strategies but to evaluate approaches for population structure control in epistasis, SNPs screened for epistasis were generated as follows. A total of only 10 candidate SNPs were selected at random from the available CEU and YRI SNP panel, with the restriction that the minor allele frequency difference between CEU and YRI was larger than *d ∈* {0.1, 0.3}. Genotypes for unlinked null loci were generated as before. A total of 1000 replicates of null data (i.e., no association between SNPs and trait) were created by repeating the process above 1000 times and by randomly assigning individuals to disease. To be able to assess power, each genetic replicate was appended with 2 functional SNPs. Disease status generation was based on 6 pure epistasis models (Supplementary material – Table S1). These models are heavily used in the epistasis field, for instance to evaluate MDR ^36^, MDR-PDT ^37^, MBMDR ^24^ and MDR-SP ^22^. They involve equal MAFs for functional SNP pairs, with MAFs ∈ {0.50, 0.25, 0.10} and no main effects. We randomly selected 2 SNPs from the pooled CEU and YRI HapMap data ensuring that the MAF in the CEU population at each of the 2 SNP was within ±0.02 of the given MAF in the chosen pure epistasis disease model. Two-locus genotypes for the functional SNP pair were then generated, conditional on fixed and equal numbers of cases and controls (100, 150 and 200) each. This process was repeated 1000 times.

#### ***Simulation setting 2***: Synthetic data derived from HapMap with structural epistasis

To introduce structural epistasis into our simulation study for GWAIS, we considered four HapMap populations: 2 closely related populations CHB and JPT (F_ST_ = 0.007) and 2 distant populations CEU and YRI (F_ST_=0.153). Then, to detect epistasis via adjusting non-linear structural differences between these populations we apply the MB-MDR methods for structured populations. We subsequently identified all significant SNP-SNP interaction pairs, adjusted for main effects and corrected for multiple testing (default options in the MB-MDR software version mbmdr-4.4.2). Based on these results, several approaches were taken to generate genotypes in the absence or presence of epistatic differences between populations. Approach 1: we generated 10,200 unlinked random genotypes including a) 10,000 SNPs randomly generated from the pooled CEU, YRI, CHB and JPT data, without association to disease and population structure, similar to simulation set 1 and b) 100 pairs of SNPs randomly selected from the aforementioned significant pairs of SNPs related to population structure comparing CHB to JPT, and CEU to YRI. From these 100 pairs, we extracted the empirical proportion of corresponding 9 two-locus genotype combinations. The associated penetrance functions were used to generate the additional 200 unlinked genotypes, by conditioning on fixed sample sizes of {100, 250} from each of the four populations. Approach 2: we simulated 110 candidate random genotypes including a) 100 without association to disease with population structure similar to Approach 1 -b) and 5 significantly interacting SNP pairs with population structure similar to Approach 1 -b). Approach 3: Two functional genotypes were randomly selected from the significant pairs that were found to be associated to population structure in such a way that the MAFs in the CEU and CHB populations at each SNP was within ±0.1 of the given MAFs in the disease model (Supplementary Table S1). A total of 1000 replicates were generated for total samples sizes of {400, 1000} and proportions of cases and controls according to 60:40.

Unlinked genotypes obtained via Approach 1 were used to extract principal components to control for population structure. The first 10 principal components were used to capture population structure in epistasis analyses. Candidate genotypes generated via Approach 2 were used to evaluate type I error rates of proposed population correction strategies in GWAIS, whereas functional genotypes as in Approach 3 were used in methods power analyses. Various ways of computing principal components were implemented to capture synthetic data underlying population structure. In particular, we considered linear principal component analysis (linear PCA), as applied to genetic data in ^23^, kernel PCA with a radial basis kernel, as implemented in the R package ***kernlab*** (Kernel-Based Machine Learning Lab), and ncMCE (non-centered multicurvilinearity embedding) kernel-based PCA introduced in ^38^ as an alternative to capture non-linear genetic differences between populations.

#### ***Simulation setting 3***: Model-based discrete populations in the absence of structural epistasis

Here, we simulated a large number of biallellic genotype frequencies for each individual in subpopulations, using Balding-Nicholas models ^39^, similar to ^34,35^. First, an ancestral allele frequency *p_a_* is randomly sampled from the uniform distribution in the interval [0.05, 0.95]. Second, Wright’s coefficient of inbreeding *F_ST_* is specified for the subpopulations *F_r_* ∈ {0.01, 0.03}, *r* = 1, 2. Third, the allele frequency 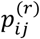 of individual *i* for genotype *j* in subpopulation *r* is simulated from a beta distribution with parameters 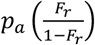 and 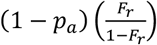, *r* = 1, 2, *i* = 1, …,*N* and *j* = 1, …,*M*. Then, genotype values {0, 1, 2} are simulated from a multinomial distribution with probabilities - computed without assuming HWE - by 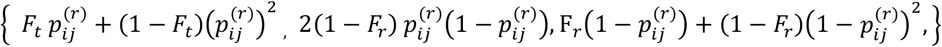 (see ^35^ and references therein). We thus generated 1000 unlinked SNPs to calculate principal components similar to ^23^. In addition, 100 SNPs were generated in a similar way as the unlinked SNPs to evaluate type I error rates. For power comparison two functional SNPs were generated taking into account the six genetic disease models presented as supplementary information (Table S1). This procedure was repeated 1000 times with total samples sizes of {500, 1000} and proportions of cases and controls according to 60:40.

#### ***Simulation setting 4***: Model-based discrete populations in the presence of structural epistasis

Instead of relying on a data-driven empirical penetrance table for structural epistasis as before (Simulation setting 2), we considered a checker-board type of model as in Table 1, which describes epistatic genetic difference between the populations using the XOR model. The parameter *β*_0_ is taken to be the average penetrance (in the absence of any genetic effect), whereas *β*_1_ captures the increase in penetrance when having the specific 2-locus genotype. In our simulations we assumed *β*_0_ = 0 and *β* = 0.35 and 0.20 for populations 1 and 2, respectively. Then, we generated 1000 unlinked random genotypes including a) 800 SNPs randomly generated similar to *Simulation setting 3* using *F_ST_* in the two subpopulations *F_r_* ∈ {0.001, 0.001}, *r* = 1, 2 ; b) 100 pairs from each population similar to *Simulation setting 2 (1b)* using the population-specific penetrance values given in Table 3 with *β*_*j*,1_ = *β*_1_ + *ε*, *j* = 1,…, 50, where *ε* is randomly drawn from *uniform* (0, 0.05). To assess type I error rate, 120 SNPs are generated of which 100 similar to (a) and 10 pairs similar to (b). A total of 1000 replicates were generated for total samples sizes of {200, 500, 1000} and proportions of cases and controls according to two scenarios 60:40 and 80:20. This dataset is used to construct principal components.

**Table 1.** Checkerboard stratification penetrance models for structural epistasis

| Population 1 |  |  |  | Population 2 |  |  |  |
| --- | --- | --- | --- | --- | --- | --- | --- |
|  | BB | Bb | Bb |  | BB | Bb | bb |
| AA | $\beta_0$ | $\beta_0 + \beta_1$ | $\beta_0$ | AA | $\beta_0 + \beta_1$ | $\beta_0$ | $\beta_0 + \beta_1$ |
| Aa | $\beta_0 + \beta_1$ | $\beta_0$ | $\beta_0 + \beta_1$ | Aa | $\beta_0$ | $\beta_0 + \beta_1$ | $\beta_0$ |
| Aa | $\beta_0$ | $\beta_0 + \beta_1$ | $\beta_0$ | aa | $\beta_0 + \beta_1$ | $\beta_0$ | $\beta_0 + \beta_1$ |

**Table 2.**
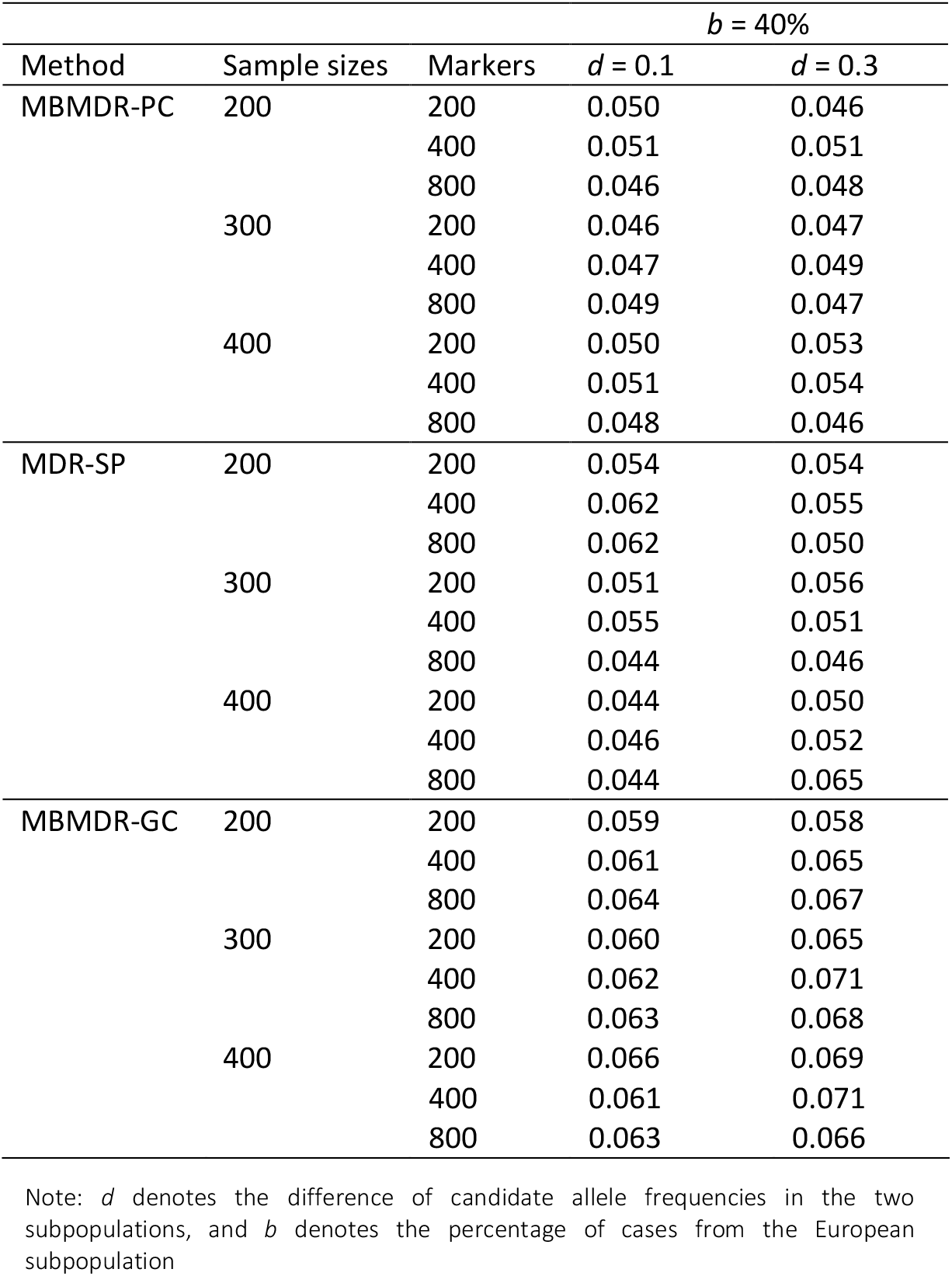
Estimates of Type I error for MBMDR-PC, MBMDR-GC and MDR-SP, with a nominal 0.05 FWER level.

**Table 3.**
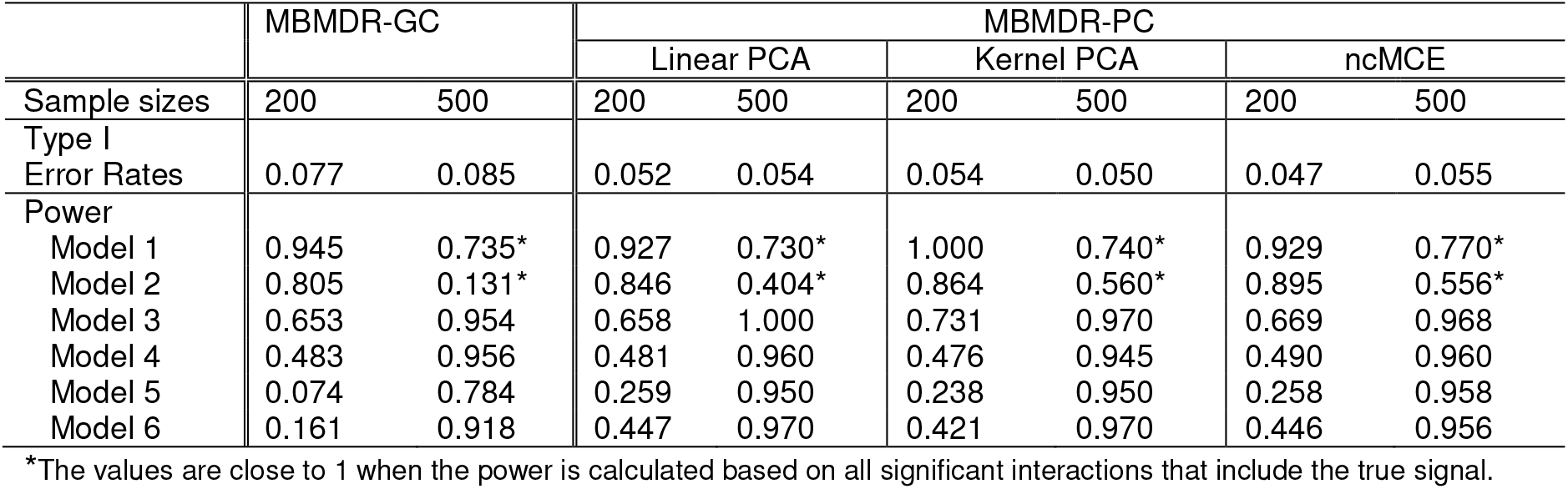
Estimates of power and type I error rates of MBMDR-PC with population structure captured by linear, kernel and ncMCE principal components and MBMDR-GC, with a nominal 0.05 FWER level.

|  | MBMDR-GC |  | MBMDR-PC |  |  |  |  |  |
| --- | --- | --- | --- | --- | --- | --- | --- | --- |
|  |  |  | Linear PCA |  | Kernel PCA |  | ncMCE |  |
| Sample sizes | 200 | 500 | 200 | 500 | 200 | 500 | 200 | 500 |
| Type I Error Rates | 0.077 | 0.085 | 0.052 | 0.054 | 0.054 | 0.050 | 0.047 | 0.055 |
| Power |  |  |  |  |  |  |  |  |
| Model 1 | 0.945 | 0.735* | 0.927 | 0.730* | 1.000 | 0.740* | 0.929 | 0.770* |
| Model 2 | 0.805 | 0.131* | 0.846 | 0.404* | 0.864 | 0.560* | 0.895 | 0.556* |
| Model 3 | 0.653 | 0.954 | 0.658 | 1.000 | 0.731 | 0.970 | 0.669 | 0.968 |
| Model 4 | 0.483 | 0.956 | 0.481 | 0.960 | 0.476 | 0.945 | 0.490 | 0.960 |
| Model 5 | 0.074 | 0.784 | 0.259 | 0.950 | 0.238 | 0.950 | 0.258 | 0.958 |
| Model 6 | 0.161 | 0.918 | 0.447 | 0.970 | 0.421 | 0.970 | 0.446 | 0.956 |

\*The values are close to 1 when the power is calculated based on all significant interactions that include the true signal.

#### ***Simulation setting 5***: Simulating genotypes for related individuals

Inspired by ^40^, we simulated 1000 replicate data consisting of 250 nuclear families, with the number of children drawn from a multinomial distribution with probabilities 1/4 to have one child, 1/2 to have two children, and 1/4 to have three children. On average, this gave rise to 1000 individuals. To generate parental genotypes, we generated 10 biallelic markers in linkage equilibrium, and assuming Hardy-Weinberg equilibrium. The allele frequencies of the functional SNP pair (*SNP*_1_, *SNP*_2_) were taken to be equal, and varied as (*p*_1_, *p*_2_) = (*p, p*), *p* ∈ (0.1,0.25,0.5). The allele frequencies of the 8 remaining non-functional SNPs were fixed at *p_j_* = 0.1 + (*j* − 3)0.05, *j* = 3,… 10. Children’s genotypes were assumed to follow Mendelian inheritance patterns. Disease penetrance for parents and children was based on Model M170, as discussed in ^41^. This epistasis model is similar to Model 1 in Table S1 (supplementary material). However, we fixed the total heritability *h*^2^ and the proportion of the total variance explained by the two-locus model variance at 0.5 and 0.05, respectively. As family relationships may induce phenotype similarity, this simulation setting was used to evaluate the performance of MBMDR-PG.

## Results

### Simulations

#### Simulation setting 1

Type I error estimates for the considered scenarios and obtained via application of MBMDR-PC, MBMDR-PG, MBMDR-GC and MDR-SP to 1000 replicated samples are presented in Table 2. In case of a single homogeneous population (CEU only) none of the estimated type I errors are significantly different from the nominal 0.05 FWER level, with a 95% confidence interval of (0.036, 0.064) ^22^. This is the case, for all considered combinations of population structure correction methods, sample sizes and number of SNPs. In case of structured samples (in particular consisting of CEU and YRI), MBMDR-PC estimated type I errors presented in Table 2 always follow Bradley’s liberal criterion. In addition, it can be seen from Table 2 that all the estimated type I error rates for MBMDR-PC are within the 95% confidence interval but it is not the case for MDR-SP. However, many type I error rate estimates based on MBMDR-GC do not fall within the 95% interval. The results of MBMDR-PG are similar to MBMDR-PC (results not shown).

In Figure 3, for allele frequencies difference *d* = 0.1 and percentage of European population *b* = 40%, MBMDR-PC is significantly more powerful than MDR-SP under all models considered in particular for small sample sizes. Moreover, MBMDR-PC outperforms MDR-SP even for large sample sizes in the models 5 and 6. Notably, these epistasis models are the toughest of the 6 considered Ritchie models in that they involve functional SNP pairs with the lowest MAFs (0.10). As the sample size increases the power of both methods increases. We also included the power results of MBMDR-PG, which are almost similar to MBMDR-PC. Similar results follow when *d* = 0.3 and *b* = 40%. In addition, the results of power based on varying number of unlinked markers are included in Figure S1 (Supplementary material) that suggest there is not much difference in the power of MBMDR-PC using 200, 400 and 800 unlinked markers for computing principal components to control population structure in our data simulation.

**Figure 3.**
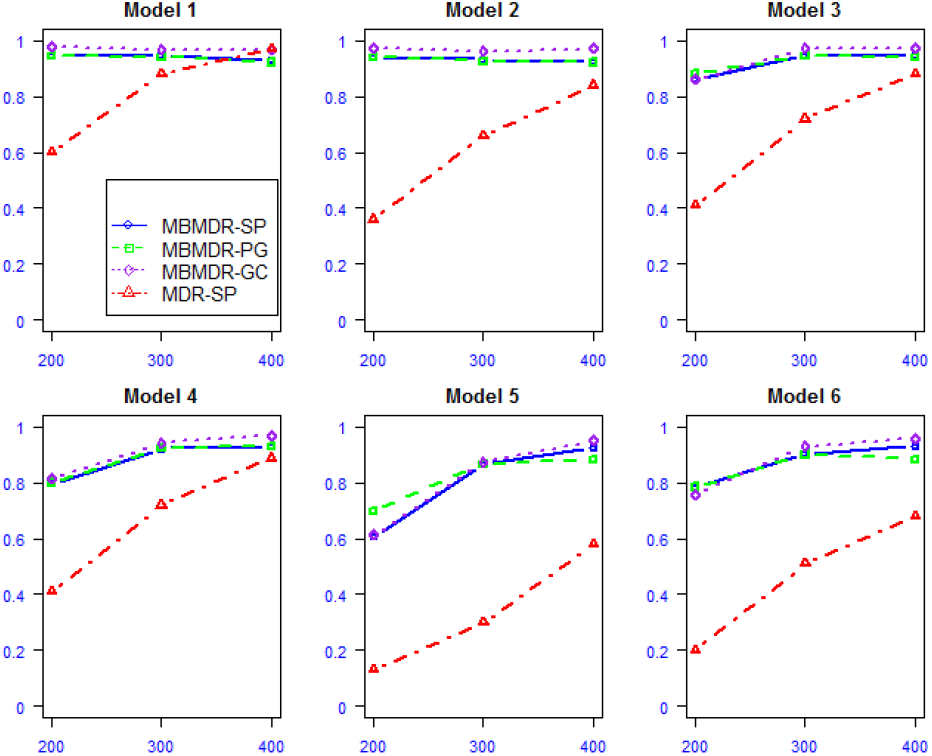
Power estimates for MBMDR-PC (blue/solid line), MBMDR-PG (green/dashed line) and MDR-SP (red/dotted line) under the six disease models based on simulated data on CEU and YRI populations with a difference of 0.3 minor allele frequency between the two populations. Percentage of cases and control from the CEU are 40% and 80%, respectively. The power (y-axis) is computed using 10 candidate SNPs. PCs are computed from 200 unlinked SNPs.

#### Simulation setting 2

The estimated power of MBMDR-PC under the discrete population simulation setting are shown in Figure 4. The results show that MBMDR-PC has high power in all scenarios of varying case-control proportions in all disease models with large samples. The power of MBMDR-PC is low for small sample sizes (100 samples from each of the two populations) for disease models with moderate and small minor allele frequencies. In general, the simulation results of varying case-control proportions have no considerable impact on the power of MBMDR-PC method for large samples of 1000 or more. The results of estimated Type I error rates for varying proportions of cases and controls with and without main effect and principal component corrections are displayed in Figure S2 (Supplementary material). From this figure we see that MBMDR-PC performs well in controlling type I error rate at the nominal 0.05 FWER level with and without main effects correction in all scenarios of case-control proportions (Figure S2 A and B). Use of the original MBMDR without population and main effect corrections in case of structured population leads to inflated type I error rates (Figure S2 D and C) in case of small samples and large difference in case-control proportions.

**Figure 4.**
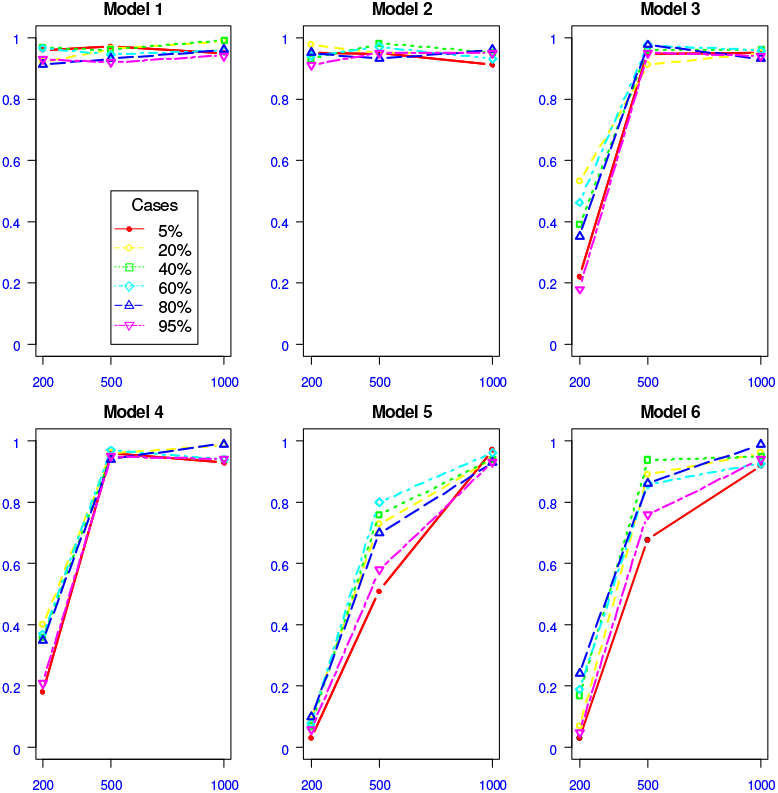
Power estimates according to varying proportions of cases and controls in six disease epistasis models and variable sample sizes (200, 500, 1000). The percentage of cases in one of the two populations are shown.

#### Simulation setting 3

To evaluate the performance of MBMDR-PC in multiple subpopulations we evaluate three principal component extraction methods: linear, kernel and ncMCE. Pairwise PC-plots for the first three principal components computed from the unlinked null SNPs are shown in Figure 5. The plot of the first and the second PCs obtained from linear PCA method (Figure 5 A1) fails to separate CHB and JPT populations. Similar result were reported in ^38^. However, the plot of the second and third linear PCs (Figure 5 A3) differentiate all four populations. In case of kernel-based PCs, the four populations are clearly separable in any of the pairwise PC plots of the first three kernel PCs (Figure 5 B1-B3). On the other hand, the plot of the first versus the second ncMCE based PCs (Figure 5 C1) was able to reveal the hierarchical structure of the four populations, reflecting the phylogenetics of these populations, as discussed in Alanis-Lobato and colleagues ^38^.

**Figure 5.**
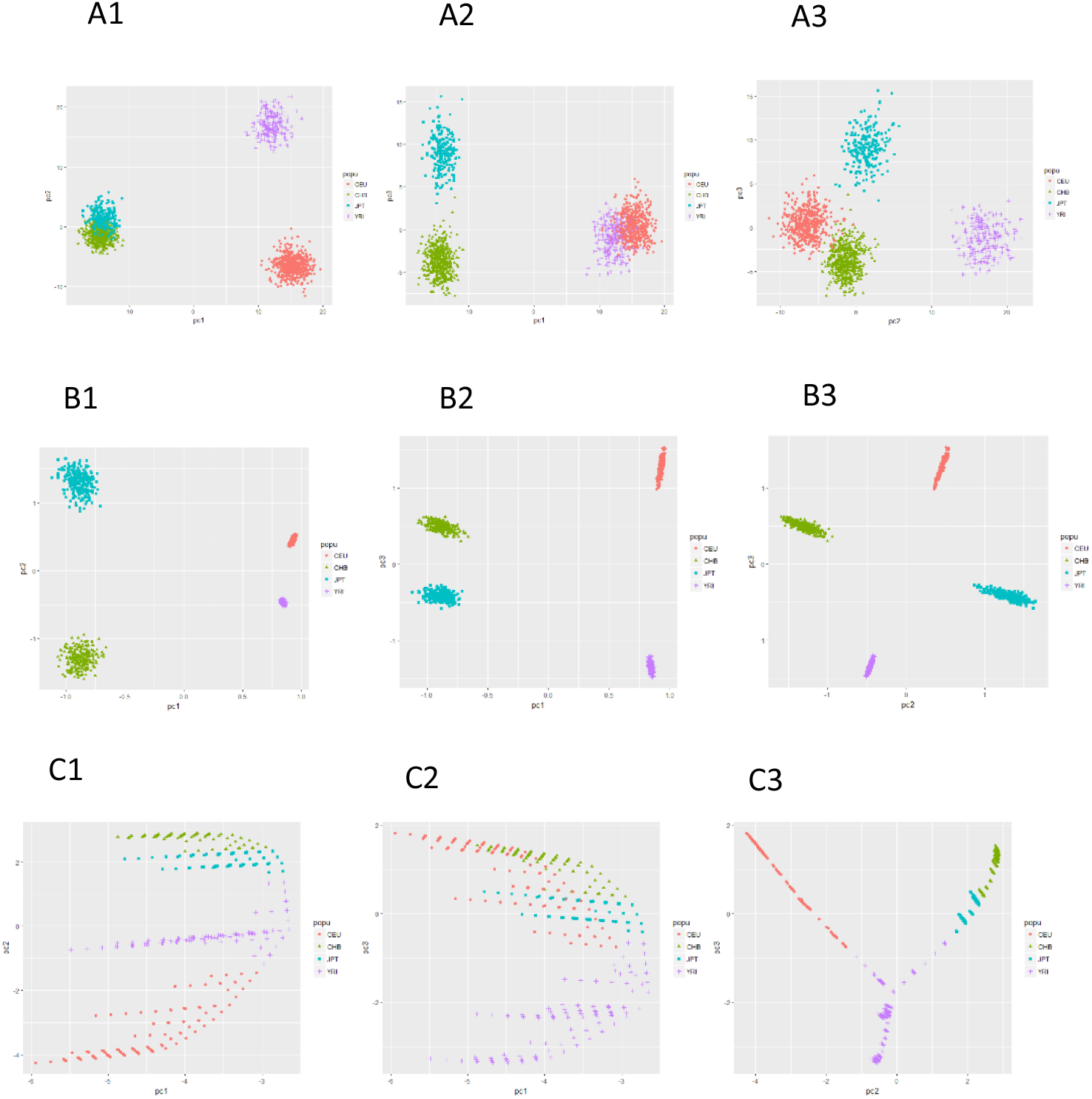
Pairwise plots of the first three principal components computed using linear (A1-A3), kernel (B1-B3) and ncMCE (C1-C3) PCA methods.

As can be seen from Table 3, none of the considered simulation scenario’s show marked differences regarding type I error control or powerto detect epistasis, when using the first 10 PCs computed via linear, kernel or ncMCE PCA methods with MBMDR-PC. In contrast, type I error estimates are somewhat inflated with MBMDR-GC. However, MBMDR-PC and MBMDR-GC give comparable power estimates, except for epistasis models with low frequency causal variants (Models 5 and 6).

#### Simulation setting 4

The scatter plot on the first 2 linear and kernel principal components for a single simulated dataset (see Methods section) is shown in Figure 6. Linear PCA indicates a non-linear genetic background structure (Figure 6 A). This is confirmed by kernel based PCA, which clearly separates the two subpopulations (Figure 6 B).

**Figure 6.**
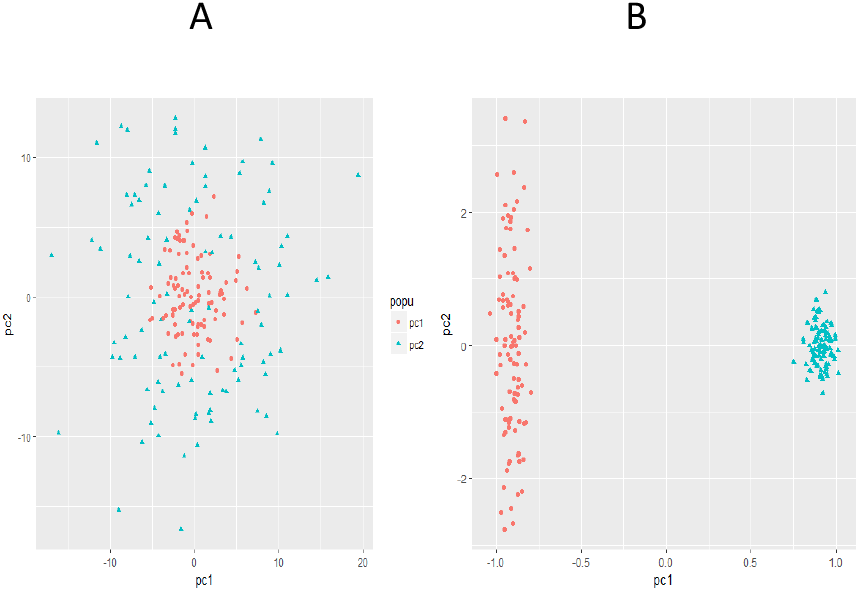
Plots of the first two principal components (A) linear PCA and (B) kernel PCA.

The estimated results of type I error rates of MBMDR-PC using linear and kernel principal components are presented in Figure 7. In the presence of phenotypic and structural epistasis, linear PCA based MBMDR-PC highly inflates the type I error which is substantially higher than the nominal 0.05 FWER level. For example, for a total sample size of 500 (cases and controls jointly) and case-control ratio is 60:40 and 80:20, the type I error rates of linear MBMDR-PC are 0.7 and 1.0 for the nominal level 0.05 (Figure 7 A and 7 B, respectively). Type I error rates of linear MBMDR-PC increases as the sample size increases. Furthermore, type I errors estimates get worse for linear PCA based MBMDR-PC with increasing levels of unbalancedness (Figure 7 B, 80:20). In comparison, the estimated type I error rates of kernel based MBMDR-PC are not significantly different from the nominal level 0.05 in all the scenarios considered.

**Figure 7.**
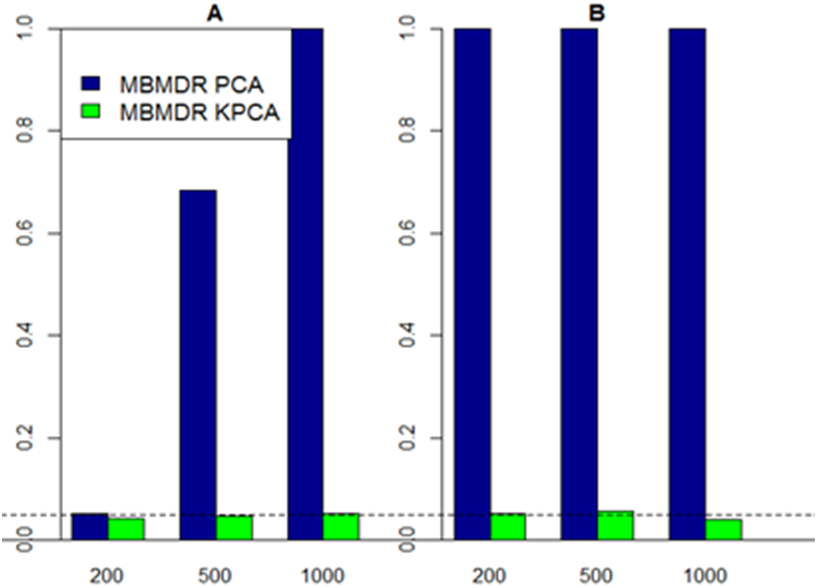
Estimated type I error rates for MBMDR-PC with case-control ratios (A) 60:40 and (B) 80:20. PC approaches considered: linear PCA (blue bars), kernel PCA (green bars).

#### Simulation setting 5

The estimated type I error rate for simulation setting based on related samples obtained from 1000 replicates (as explained in the Methods section) is 0.051., which is close to the nominal 0.05 level. Power estimates for epistasis model M170 (see Methods) increase with increasing minor allele frequencies for the causal epistasis SNP pair (0.45, 0.885 and 0.911 for MAFs of 0.1, 0.25 and 0.5, respectively).

## Discussion

In GWAS, Structured Association (SA) ^42–45^, Genomic Control (GC) ^34,35^, Principal Component Analysis (PCA) ^23^ and Mixed Modeling (MM) ^46^ are the main 4 strategies to deal with confounding associations due to shared genetic ancestry. The basic idea of SA is to infer the underlying population structure and then to incorporate this information in subsequent testing for genetic associations of interest. In contrast, the basic idea of GC is to correct the null distribution of genetic association tests for the effects of the unspecified population structure ^34^. The statistical advantage of SA methods depends on the degree of information provided by the available marker data to make inferences about the true structure. GC methods rely on adjusting all marker-trait associations in the same way, which ignores the strength of the relationship between the genealogy of the genetic marker under study and the (hidden) pedigree structure, and thus also dependencies between markers. The basic idea of mixed models (MM) for controlling population structure is to account for pairwise relatedness between individuals, for example, using a kinship matrix. It is an approach that naturally accommodates familial and cryptic relatedness in the data. Since mixed models have long been computationally too intensive it took until the development of more efficient algorithms for them to gain popularity again. Some of the algorithm improvements are incorporated in the following approaches: compressed-MLM ^46^, EMMA (Efficient Mixed-Model Association) ^47^, EMMAX (EMMA eXpedited) ^48^, GEMMA (Genome-Wide Efficient Mixed-Model Association) ^49^, LRLMM (low rank linear mixed model ^32^, FaST-LMM (Factored Spectrally Transformed Linear Mixed Model) ^50^, FaST-LMM-Set ^51^, GRAMMAR-Gamma (fast variance components-based two-step method) ^52^, and FarmCPU (Fixed and random model Circulating Probability Unification) ^53^. Principal Components Analysis (PCA) allows data transformation to a new coordinate system such that the projection of the data along the first new coordinate has the largest variance, the second principal component has the second largest variance, and so on. The relative straightforwardness of PCA, its ease of use, the availability of efficient algorithms and its ability to detect individuals with unusual or differential ancestry ^54–59^ has made PCA among the most heavily used strategies in the context of genetic association studies in structured populations.

Once principal components are obtained, several choices can be made to use these for the purpose of confounding correction in GWAS. Assuming that the GWAS is performed within a regression framework, the most straightforward approach is to include the first few principal components, capturing genetic ancestry of each individual, as fixed effects in a (generalized) linear model. Alternatively, instead of directly including the principal components in a regression model, both phenotype and genotypes can be adjusted by top PCs as in EIGENSTRAT ^23^. The adjusted phenotype is defined as the residual of fitting a linear regression model of phenotype on a number of principal components. A similar model fitting is performed to obtain adjusted genotypes. The idea is to remove the confounding effect due to population structure from both phenotype and genotypes, as should be. In case-control studies, where the phenotype is encoded as a binary variable, logistic regression rather than linear regression can be employed to obtain adjusted phenotypes. In general, depending on the nature of variables under consideration, whether at the phenotype or genotype level, appropriate generalized linear models can be considered to obtain population structure adjusted phenotypes and genotypes ^12^.

The aforementioned methods naturally extend to epistasis detection frameworks, in particular those that allow for a regression model component in their methodology. One such framework is MB-MDR (32), which adds a model-based component to Multifactor Dimensionality Reduction to adjust for lower order effects or confounders (46). Our proposed MBMDR-PC, MBMDR-PG and MBMDR-GC methods for detecting epistasis in the presence of population structure are built on the Model-Based Multifactor Dimensionality Reduction (MB-MDR) method, as described earlier ^10,24,25^. MBMDR-PC and MBMDR-PG involve first deriving adjusted phenotypes (residuals) obtained from fitting logistic and mixed logistic regression models with the original genotypes. The first few principal components (linear or non-linear) are used as covariates for fitting logistic regression and the kinship matrix is used to determine the covariance structure of the random effects.

Population structure in GWAS is usually simulated using relatively simple models, largely ignoring dependencies between ancestry informative markers beyond LD, such as structural epistasis. In this work, we not only considered epistatic ancestry informative, but also paid special attention to the idea of non-linearity in population genetics ^38^. We have also considered the impact of unequal sample sizes on GWAIS for structured populations.

In this work, we have highlighted the importance of detecting and correcting population structure in epistasis studies. It has been established that not accounting for population structure due to allele frequency differences among populations and subpopulations can result in high false-positive results or reduced power in genome-wide association studies. Similarly, in epistasis studies, using extensive simulation settings we have shown that failure to account for complex population structure results in inflated false positives or low power to detect true signals of epistasis.

In general, our simulation results show that in the presence of population structure MBMDR-PC and MBMDR-PG consistently control type I error rate at the nominal level compared to MBMDR-GC which has a slightly inflated type I error rate. Our three methods of population structure correction are more powerful than MDR-SP. Thus, MBMDR-PC and MBMDR-PG for GWAIS adjusted for confounding by (non-)linear population structure give promising results and are to be preferred over MDR-SP in the considered simulation settings. For related samples, MBMDR-PG based on a generalized linear mixed model should be used. To minimize spurious interaction signals induced by strong main effects, a codominant correction for main effects is advised. In many instances of mild population structure, MB-MDR with codominant correction for lower order effects has a comparable performance to MBMDR-PC. All analyses can easily accommodate covariates using similar principles as in MBMBDR-PC and MBMDR-GC.

The outperformance of MBMDR-PC clearly depends on the ability of the principal components to capture population structure well. Our choice of the checker board stratification model is to inject strong non-linearity in the genotype data but we believe that more work is required in defining stratification models. In such more complex non-linear population structure, the widely used linear principal component analysis fails to properly differentiate populations. Alternatively, kernel based PCA often differentiate non-linear population structure. Our simulation result that compares MBMDR-PC based on linear and kernel principal components shows that MBMDR-PC with linear principal components has inflated type I error and become worse as the sample size increases and the ratio of case-control becomes more unbalanced. On the contrary, MBMDR-PC based on kernel principal components effectively control population structure and maintain the type I error rate at the required nominal level. Thus, not properly accounting for complex genetic substructures in GWAIS dramatically increases false positive epistasis findings and kernel-based strategies in GWAIS deserve more attention. But then, efficient computational tools to extract non-linear PCs from large consortium genetic data sets would be welcome.

In conclusion, MBMDR-PC is a generally well performing approach, compared to the computationally intensive MBMDR-PG and MBMDR-GC approaches, although its performance is highly depending on how well PCs capture population structure. We recommend to use both linear and nonlinear versions of PCA, whenever possible. MB-MDR analytics should be seen as part of an entire analysis pipeline which involves making marker selection choices and performing post-analysis steps to validate and replicate findings, as well as to seek biological evidence for flagged interacting regions with MB-MDR ^18^. Fast implementation for multiple testing correction in exhaustive epistasis screenings ^10^ makes these MB-MDR based methods efficient tools for GWAIS in structured populations. Our work is important in view of ongoing initiatives of epistasis detection in large-scale heterogeneous consortium data, as we have shown that inadequate capturing of population structure is devastating in GWAIS for obtaining meaningful and replicable epistasis results.

## Supplemental Data

Supplemental data include additional materials on methods and one table and two figures from simulation studies.

## Acknowledgement

We thank Myriam Nemry for suggestions regarding the implementation of ideas in the MBMDR software. This research was in part funded by the Fonds de la Recherche Scientifique (F.N.R.S.), in particular “Integrated complex traits epistasis kit” (Convention n° 2.4609.11) [KVS]. We also acknowledge research opportunities offered by F.N.R.S., including “Foresting in Integromics Inference” (Convention n° T.0180.13) [KC], and by the interuniversity research institute Walloon Excellence in Lifesciences and BIOtechnology (WELBIO) [FA, KVS].

## Web Resourses

MBMDR-PC, MBMDR-PG and MBMDR-GC are available via the MBMDR software (from version mbmdr-4.4.1 onwards), which is downloadable from http://bio3.giga.ulg.ac.be/index.php/software/mb-mdr/. Main options

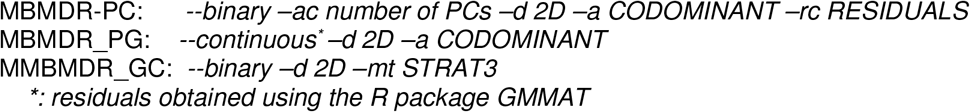

