## Supplemental Data for "Epistasis Detection using Model Based Multifactor Dimensionality Reduction in Structured Populations"

### Supplementary Material to Epistasis Detection using MBMDR in Structured Populations

Fentaw Abegaz[1,*], François Van Lishout[1], Jestinah M Mahachie John[1], Kridsadakorn Chiachoompu[1], Archana Bhardwaj[1] , Elena S Gusareva[1], Zhi Wei PhD[2], Hakon Hakonarson[3,4], and Kristel Van Steen[1,5], on behalf of the International IBD Genetics Consortium

[1] GIGA-R, Medical Genomics – BIO3, University of Liege, Liege, Belgium
[2] Department of Computer Science, New Jersey Institute of Technology, Newark, NJ, USA
[3] Center for Applied Genomics, The Children's Hospital of Philadelphia, Philadelphia, PA, USA
[4] Division of Human Genetics, Department of Pediatrics, The Perelman School of Medicine, University of Pennsylvania, Philadelphia, PA, USA
[5] WELBIO (Walloon Excellence in Lifesciences and Biotechnology), University of Liege, Liege, Belgium

#### Method and Materials

##### Principal Components Analysis (PCA) in the absence of family relatedness

The popular EIGENSTRAT software to correct for population structure in GWAS contexts uses top linear PCs as covariates in a multiple regression [1]. Many ad hoc procedures and formal statistical tests exist to determine the number of principal components to correct for population structure. It is a common practice to take between 2-10 principal components for correcting population structure in genome-wide association studies involving several countries. Even though linear PCA is most popular and adequate in most cases to capture ancestry genetic background, PCA may fail to capture non-linear population structure in genetics as shown in [2]. The non-linear method developed by Alanis-Lobato and colleagues is based on a non-centered Minimum Curvilinear Embedding (ncMCE) kernel. Whereas the latter is able to better capture phylogenetic signals in samples, PCA better seems to reflect geographic dependencies [3]. Alternatively, kernel-based PCA can be adopted to account for non-linear structures in high dimensional genetics data.

##### Linear PCA

In genome-wide association studies, principal component analysis has been used to explicitly model ancestry differences between cases and controls along continuous axes of variation [1]. The computation of principal components is as follows. Suppose there are $N$ individuals genotyped at $M$ SNPs. For each individual $i$, let $\mathrm{G}_{ij}$ be the genotype data taking a value 0, 1 or 2, that represents the number of copies of minor alleles and let $Y_i$ be the phenotype information. For the jth SNP, the mean is

$$\mu_j = \frac{\sum_{i=1}^{N} G_{ij}}{N}$$

and the standard deviation is

$$\sigma_j = \sqrt{p_j(1-p_j)}$$

where $p_j$ is given by either as $p_j = \left(1 + \sum_i z_{ij}\right)/(2+2N)$ or $p_j = \mu_j/2$ and defined as a posterior estimate of the unobserved underlying allele frequency or an estimate of the underlying allele frequency of SNP $j$, respectively.

The classical linear PCA relies on finding the eigenvectors of the matrix $X$ or the $N \times N$ covariance matrix $XX^T$ or the $M \times M$ covariance matrix $X^TX$ (for convenience we ignore the factor $1/N-1$). In genetics where $M \gg N$, matrix decomposition on $XX^T$ has computational advantage over $X^TX$. The eigenvectors or principal axis of variations can be obtained by using either singular value decomposition of the matrix X

$$X = U\mathrm{D}V^T$$

or the eigen-decomposition of the marix $XX^T$

$$XX^T = U\Lambda U^T,$$

where $U$ is a matrix of eigenvectors and $diag(\Lambda) = \lambda_1, \ \ldots \ , \ \lambda_r = diag(D^2)$ are the eigenvalues. Then, the principal components of the data are given by the projection of the data onto the eigenvectors

$$W = XV = U\Lambda^{1/2}.$$

**Kernel PCA**

The procedure to extract the kernel PCs is as follows as presented in [4]. The first step in kernel PCA is mapping $N \times M$ dimensional input genotype data $G$ into a higher dimensional feature space using non-linear transformation function of $\Phi(G)$. Then, using the inner products of the new feature vectors to form a kernel matrix $K$ of dimension $N \times N$ with the $(jk)$-th entry equals $K(G_j, G_k) = \Phi(G_j)^T\Phi(G_k)$. However, the mapping of original data to a very high-dimensional feature space makes it difficult to compute the inner product. This problem can be circumvent in some cases by using the "kernel trick". The kernel trick allows one to efficiently compute the kernel $K(G_j, G_k)$ using a kernel function which depends on the dimensionality of the original data points $G_j$ and $G_k$ but avoids explicit mapping of the data points $G_j$ and $G_k$ to the higher-dimensional $\Phi(G_j)$ and $\Phi(G_k)$. For example, using the radial basis function (RBF) the Kernel is computed as

$$K(G_j, G_k) = \frac{\exp\left(\left\|G_j - G_k\right\|^2\right)}{2\sigma^2} = \Phi(G_j)^T\Phi(G_k)$$

Note that since centered data is required to perform an effective principal component analysis, a centered kernel $K_c$ is obtained by

$$K_c = K - 1_N K - K 1_N + 1_N K 1_N$$

where $1_N$ denotes a $N \times N$ matrix for which each element takes value $1/N$. The kernel based principal components can be obtain using eigenvalue decomposition of the centered kernel

$$K_c = U^{(K)} \Lambda^{(K)} {U^{(K)}}^T$$

where $U^{(K)}$ is a matrix of eigenvectors and $\Lambda^{(K)}$ is a diagonal matrix of positive eigenvalues of $K_c$. It follows that the $j$th kernel principal component score corresponding to $j$th eigenvector based on the centered kernel is given by

$$W_j^{(K)} = \sum_{i=1}^{N} U_{ij}^{(K)} \, \mathrm{K}_c(Z_i, Z), j = 1, \dots, N.$$

It is a common practice to take the first 5 or 10 principal components for correcting population structure in association studies. There are also many ad hoc procedures and formal statistical tests to determine the number of principal components to correct for population structure.

Table S1: Pure epistasis disease models used in the simulation to evaluate power of MBMDR methods for structured populations

Model 1, $p = 0.5$

| | BB | Bb | bb |
|---|---|---|---|
| AA | 0 | 0.1 | 0 |
| Aa | 0.1 | 0 | 0.1 |
| aa | 0 | 0.1 | 0 |

Model 3, $p = 0.25$

| | BB | Bb | bb |
|---|---|---|---|
| AA | 0.08 | 0.07 | 0.05 |
| Aa | 0.1 | 0 | 0.1 |
| aa | 0.03 | 0.1 | 0.04 |

Model 5, $p = 0.1$

| | BB | Bb | bb |
|---|---|---|---|
| AA | 0.07 | 0.05 | 0.02 |
| Aa | 0.05 | 0.09 | 0.01 |
| aa | 0.02 | 0.01 | 0.03 |

Model 2, $p = 0.5$

| | BB | Bb | bb |
|---|---|---|---|
| AA | 0 | 0 | 0.1 |
| Aa | 0 | 0.05 | 0 |
| aa | 0.1 | 0 | 0 |

Model 4, $p = 0.25$

| | BB | Bb | bb |
|---|---|---|---|
| AA | 0 | 0.01 | 0.09 |
| Aa | 0.04 | 0.01 | 0.08 |
| aa | 0.07 | 0.09 | 0.03 |

Model 6, $p = 0.1$

| | BB | Bb | Bb |
|---|---|---|---|
| AA | 0.09 | 0.001 | 0.02 |
| Aa | 0.08 | 0.07 | 0.005 |
| aa | 0.003 | 0.007 | 0.02 |

#### Results

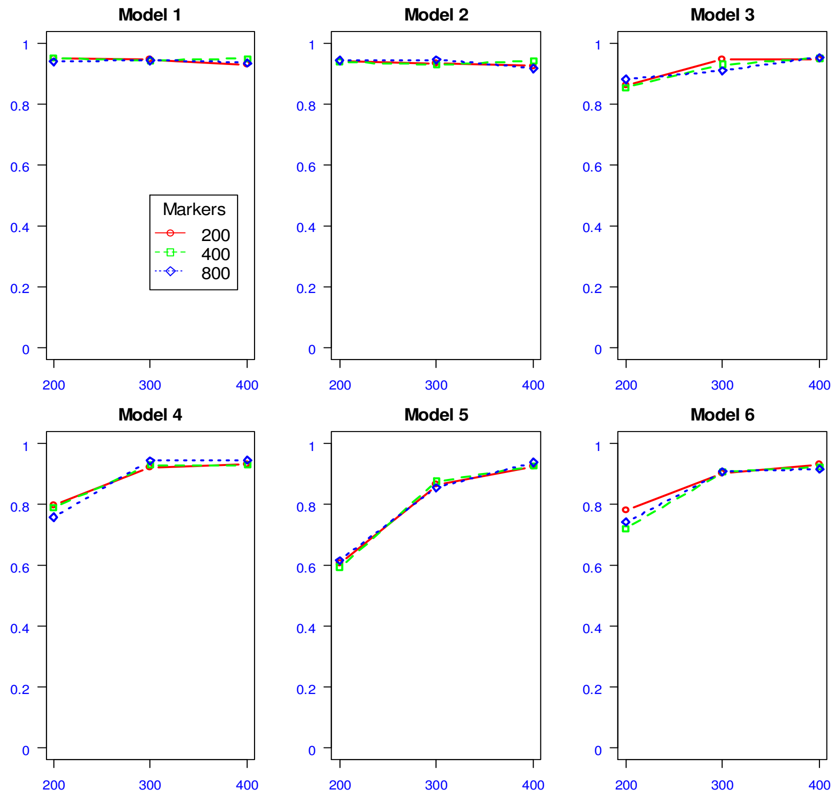


Figure S1. Power comparisons of MBMDR-PC of varying number of SNPs in PC computation under the six disease models based on simulated data on CEU and YRI populations with a difference of minor allele frequencies of candidate SNPs greater than 0.3 between the two populations and percentage of cases and control from the CEU are 40% and 80%, respectively. The power (y-axis) is computed using 10 candidate SNPs and 200 (red), 400 (green) and 800 (blue) unlinked SNPs to control population structure via principal components with varying sample sizes (x-axis).

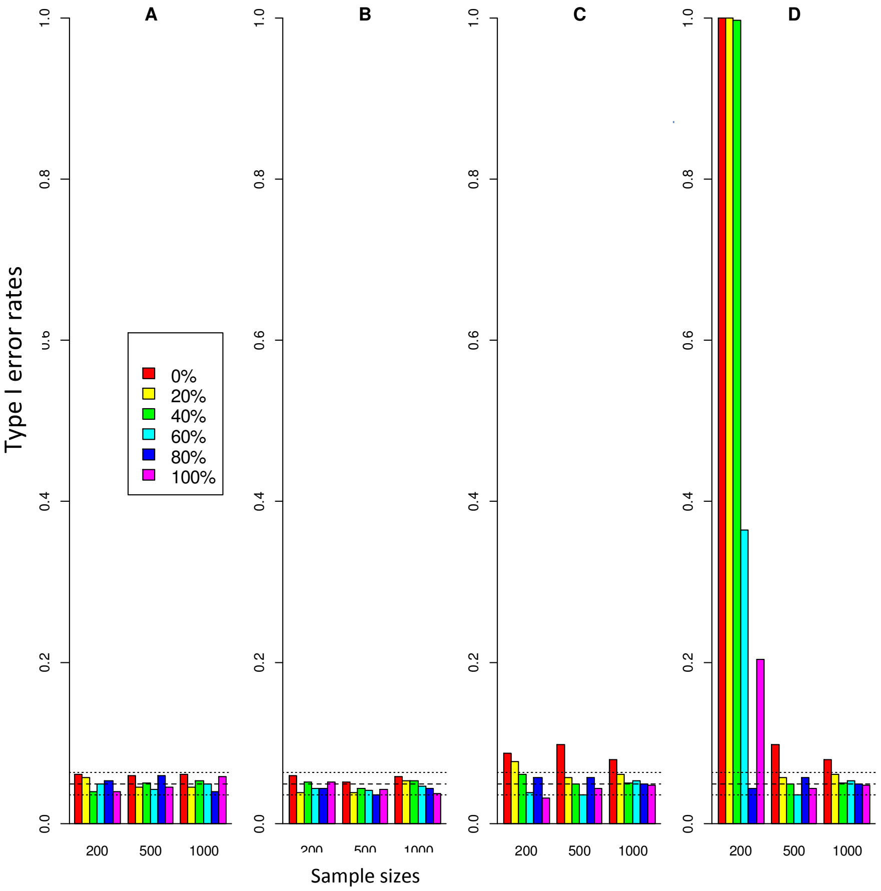


Figure S2. Results of type I error according to varying proportions of cases and controls. The percentage of cases in one of the two populations are shown. MBMDR methodology with (A) PCs and main effects correction, (B) only PCs correction, (C) only main effects correction, and (D) no correction.
